# Structural insights into plasticity and discovery of remdesivir metabolite GS-441524 binding in SARS-CoV-2 macrodomain

**DOI:** 10.1101/2021.03.04.433966

**Authors:** Xiaomin Ni, Martin Schröder, Vincent Olieric, May E. Sharpe, Victor Olmos, Ewgenij Proschak, Daniel Merk, Stefan Knapp, Apirat Chaikuad

## Abstract

The nsP3 macrodomain is a conserved protein interaction module that plays essential regulatory roles in host immune response by recognizing and removing posttranslational ADP-ribosylation sites during SARS-CoV-2 infection. Thus, targeting this protein domain may offer a therapeutic strategy to combat the current and future virus pandemics. To assist inhibitor development efforts, we report here a comprehensive set of macrodomain crystal structures complexed with diverse naturally-occurring nucleotides, small molecules as well as nucleotide analogues including GS-441524 and its phosphorylated analogue, active metabolites of remdesivir. The presented data strengthen our understanding of the SARS-CoV-2 macrodomain structural plasticity and it provides chemical starting points for future inhibitor development.

---

The recently emerge coronavirus SARS-CoV-2 has become a major global health concern. SARS-Cov-2 is an enveloped (+) single-stranded RNA ((+)ssRNA) coronavirus, encoding several functionally relevant enzymes and protein interaction domains that are necessary for the viral life cycles and deregulation of the innate immune response of the host^1-2^. Non-structural protein 3 (nsP3) is one of the most complex coronavirus proteins comprising several domains and activities^1, 3-5^. The coding sequence includes a macrodomain (MD) module, which is structurally conserved in coronaviruses and generally binds ADP-ribose^6^. The viral MDs have been shown to play a pivotal role in recognizing and removing posttranslational ADP-ribosylation from specific sites ^6-9^. This posttranslational modification, which may occur as poly-ADP-ribosylation or mono-ADP-ribosylation (MARylation), is catalyzed by PARP/ARTD enzymes and it is associated with stress signaling and innate immune response against pathogens^10-11^. Several MARylating PARP enzymes are activated in response to pathogen-associated molecular patterns and interferon^9, 11^, suggesting therefore that MARylation is a key modification of the innate immune response against viruses.

Several studies have provided strong evidence that viral MDs, including those of several coronaviruses, such as SARS-CoV and MERS-CoV, and different alphaviruses, such as Chikungunya virus and O’nyong’nyong virus, function as hydrolases capable of removing ADP-ribose from proteins ^6, 9, 12-14^. Similarly, such de-mono-ADP-ribosylating activity (de-MARylation) have been recently demonstrated for the first nsP3 macrodomain, known also as the Mac1, of SARS-CoV-2^15-16^, while the other two MDs likely exert distinct activities^6^. The macrodomain and its de-MARylation function is important for efficient viral replication as viruses harboring inactive MDs exhibit reduced replication ability and pathogenicity^10, 17^ and are sensitive to interferon pretreatment^18-19^. In addition, de-MARylation activity of the viral macrodomain has been convincingly linked directly or indirectly to the function of viral proteins involved in viral genome replication and/or in antagonizing the innate immune response^9-10^.

The central role of the SARS-CoV-2 macrodomain in viral replication and in mediating inflammatory responses in the host suggests that targeting this protein module may offer a therapeutic strategy for combating severe respiratory disease. Recent studies have elucidated the crystal structure of this protein module^15-16, 20-22^, demonstrating that the macrodomain of SARS-CoV-2 exhibits macro-H2A topology highly similar to homologues of other viruses including SARS-CoV^9, 16, 23-24^. The pocket is lined with a set of highly conserved amino acids and catalytic residues, thus its interaction with ADP-ribose resembles that of other viral macrodomain homologues^16, 20^. Nevertheless, some subtle structural differences have been observed, suggesting that SARS-CoV-2 macrodomain may possess some degree of plasticity^20^, opening a possibility for binding of diverse small molecules^15^.

Thus, to strengthen our understanding on the nature and druggability of SARS-CoV-2 macrodomain we reported a collective set of crystal structures, including an apo state as well as high resolution structures of complexes with diverse naturally-occurring nucleosides and nucleotides. Comparison of these structures highlighted ligand induced domain plasticity of the ADP-ribose binding site. In addition, we described an unprecedented discovery of the binding of GS-441524 and its phosphorylated analogue, active remdesivir metabolites.

In order to understand plasticity of the macrodomain we first attempted to crystallize an apo state, and obtained two orthorhombic crystals at 2.16-2.20-Å resolution. The presence of five and three molecules in their asymmetric units suggested therefore potential flexibility of the protein. Indeed, structural comparison of all chains revealed that while the core β-sheet and helices superimposed well, some differences in the conformations of residues lining the ADP-ribose binding pocket were evident (Figure 1A and 1B). By comparison, the AMP binding site was more rigid than the ribose-1-phosphate binding site, in which all three loops connecting β3-α2, β5-α4 and β6-α5 that lined the pocket exhibited various degrees of flexibility. Similar to previous report^20^, we observed diverse conformations of Phe132 and Ile131 located in the phosphate binding site (Figure 1B). The flexibility of the phenylalanine and isoleucine was rather interesting as it potentially determines the availability of the pocket to diverse ligands. This role was supported by conformational changes observed in the complexes with MES and HEPES molecules that were used as buffers in crystallization reagents and occupied the ribose-1-phosphate binding site (Figure 1C and 1D). Both buffer molecules bound within the phosphate binding side similarly to previous observation^20^, and caused conformational changes of loop β6-α5 residues, in essence an out-swing of Phe132 and a rotation of Ile131 side chain. In comparison, slight difference was observed for the binding of MES that induced also Ala129-Gly130 main chain flip, enabling two hydrogen bonds which were absent in the HEPES complex (Figure 1C and 1D). Overall, the structural changes seen in the apo state and both complexes with these weakly interacting ligands suggested by their high B-factors revealed therefore potential intrinsic plasticity of the macrodomain especially within its ribose-phosphate binding pocket, which may form a determinant factor for the binding of ligands.

**Figure 1.**
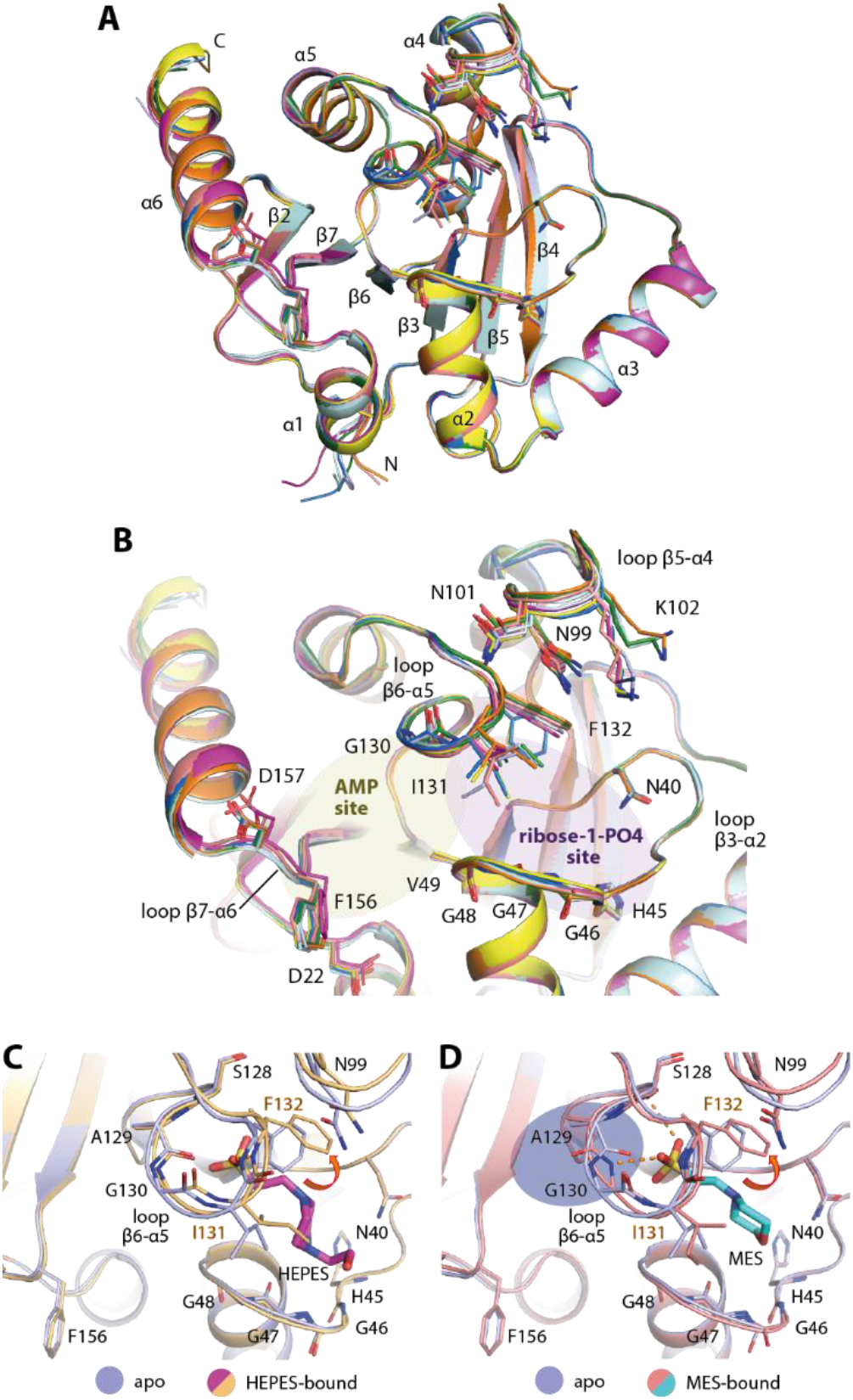
Plasticity of the ADP-ribose binding pocket of SARS-CoV-2 macrodomain. A) Superimposition of eight molecules from the asymmetric units of two distinct apo crystal structures (pdb id 6ywk and 6ywm). B) Closed up of the ADP-ribose binding pocket of all eight molecules reveals flexibility of the residues lining the pocket, notably Phe132 and Ile131. C, D) Binding of HEPES (pdb id 6ywk) and MES (pdb id 6ywm) used as crystallization reagents within the ribose-phosphate biding site requires conformational changes of Phe132 and Ile131 necessary for an accommodation of the sulfate moieties of the ligands.

To further understand the conformational changes necessary for the binding of ADP-ribose, we performed soaking of the ligand into the apo crystals, and compared the ligand-bound form (2.50-Å resolution) with the apo state. Interestingly, a number of significant side chain alterations were observed primarily at two regions: loop β6-α5 as well as loop β3-α2 that constructed the ribose-phosphate binding site (Figure 2A). Substantial alterations were noted for the former structural motif, which included an ∼56° outswing of Phe132 vacating the phosphate binding pocket in addition to a rotation of Ile131 side chain to pack on top of the terminal ribose and a main chain carbonyl flip of A129 to enable hydrogen bond interactions between Gly130 amine and the phosphate moiety of the ADP-ribose. At the opposing site of the binding pocket, small alterations of the glycine-rich loop β3-α2 were also evident, involving the rearrangement of Gly46-47 main chains bringing the loop closer to the ligand. Nevertheless, two configurations of this glycine-rich loop were observed for the ADP-ribose complex; one had Gly47 carbonyl atom flipped ‘in’ (two of five monomers) and the other conformation with an ‘out’ oriented carbonyl group (in the other three monomers; Figure 2B and 2C). These structural changes provided further evidence for high flexibility of this glycine-rich loop, which contains several catalytic residues important for enzymatic function^16^. Nonetheless, the two conformations of the loop did not alter the contacts between the terminal ribose and the protein, which were also highly conserved compared to similar complexes that have been reported previously^16, 20^.

**Figure 2.**
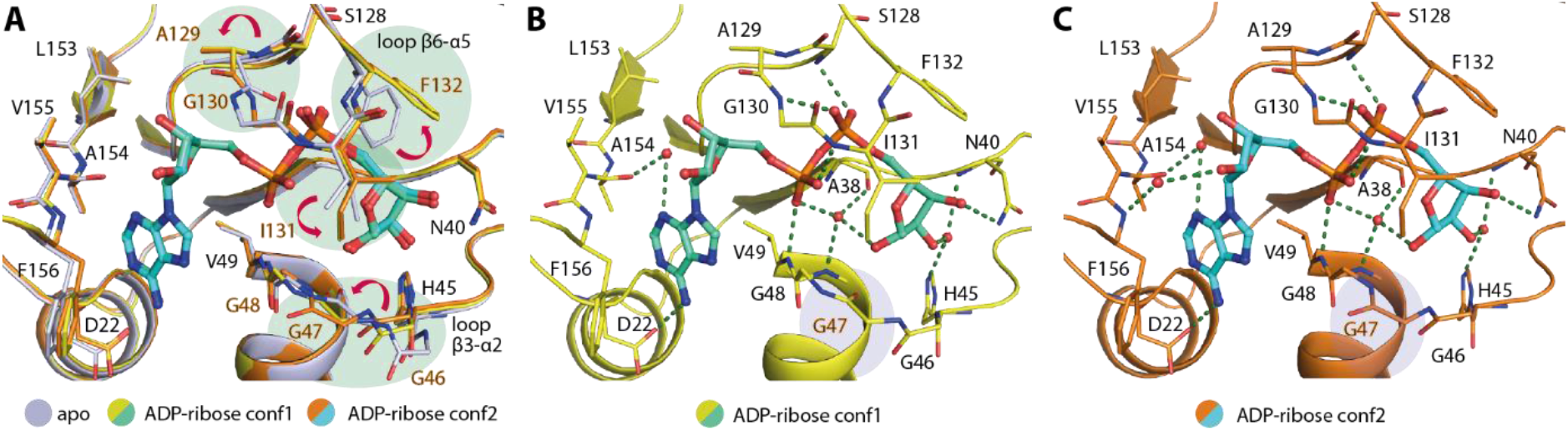
Binding of ADP-ribose triggers conformational changes. A) Superimposition of the apo form and two conformations of the ADP-ribose complex (conf1 and conf2; pdb id 6ywl) reveals several conformational changes potentially necessary for the binding of the ligand. B) and C) Two conformations observed for the ADP-ribose complex with a notable difference in the configurations of Gly47 backbone. Water molecules are shown as red spheres and green dashed lines indicate hydrogen bonds.

Plasticity of the ADP-ribose binding pocket of the macrodomain prompted us to question whether it might be able to accommodate other nucleosides and nucleotides. Thus, we attempted to co-crystallize the protein with selected naturally-occurring ligands, including AMP, GDP-glucose, ribose-1-phosphate, β-NAD and β-NADP, and all of which except ribose-1-phosphate led to successful determination of complexed structures at 1.55-2.05-Å resolution. Examination of the electron density suggested however modification of the bound ligands (Figure 3). In the case of AMP, only the adenosine moiety, the product of autohydrolysis lacking the phosphate group, was presented in the structure. Interestingly, the adenosine ligand exerted two binding modes; one resembled that of the adenosine moiety in the ADP-ribose complex, whereas a slight tilt of the adenine ring in the second binding mode positioned the ribose group outside in the solvent exposed region (Figure 3A and 3B). Although the hydrogen bond between the adenine amine group and Asp22 was still maintained, different water-mediated contacts between the ribose moiety and the protein were evident.

**Figure 3.**
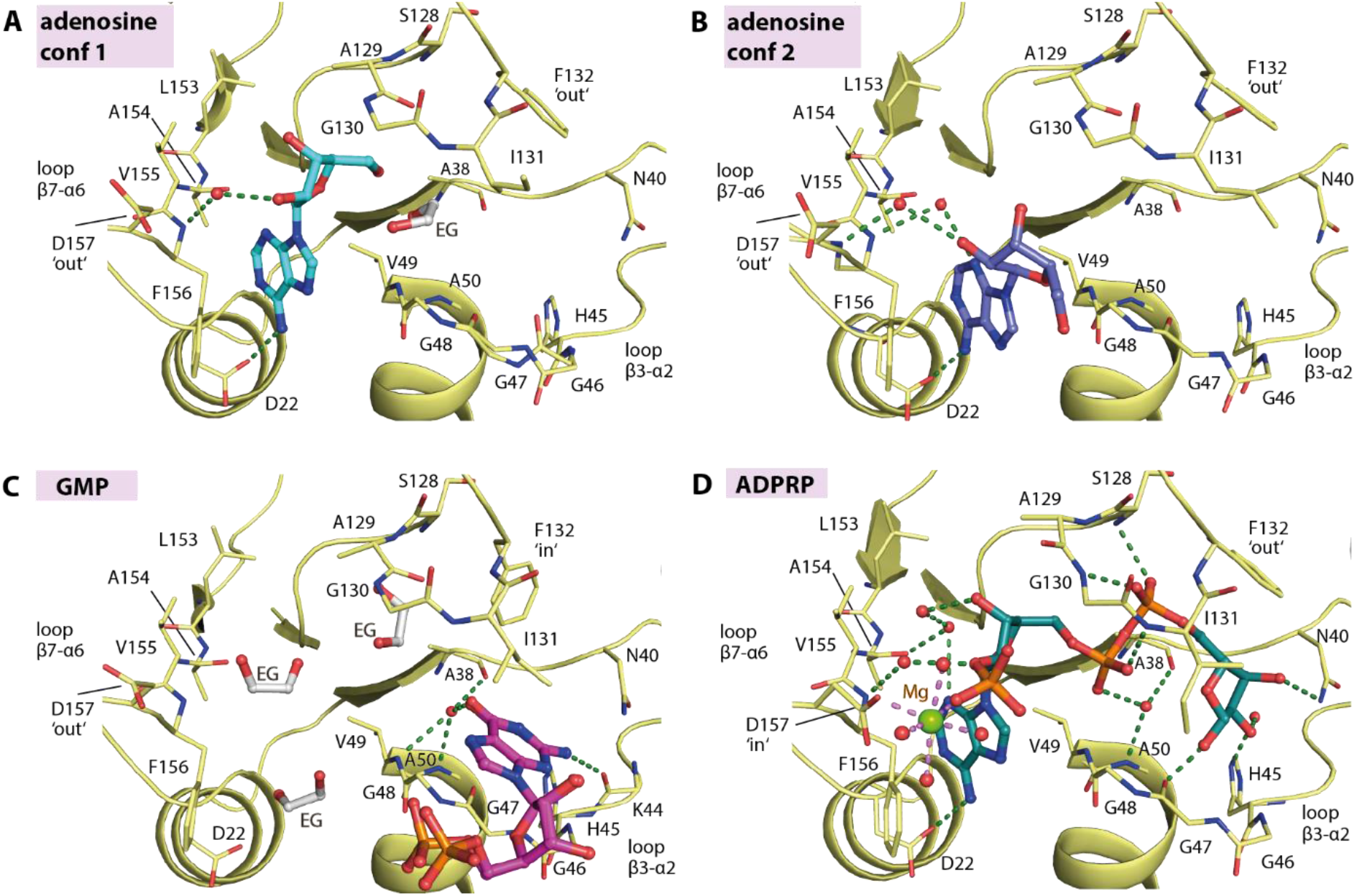
Nucleoside and nucleotide binding in the macrodomain. The binding of adenosine in two different conformations (A-B; pdb id 7bf3), GMP (C; pdb id 7bf4) and ADP-ribose-2’-phosphate (ADPRP; D; pdb id 7bf5). The ligands are shown as stick representation with water molecules shown as red spheres and ethylene glycol molecules labelled as EG.

For GDP-glucose, the electron density only allowed modelling of GMP, a product of hydrolysis. Nonetheless, the binding of GMP in the ribose-1-phosphate binding pocket of the macro-domain was intriguing (Figure 3C). The guanine ring surprisingly occupied the cavity reserved for the terminal ribose moiety of ADP-ribose, not the AMP-binding site likely due to a poor fit between the guanine 6-hydroxyl and 2-amine groups and Asp22 and the backbone amide of Val155 and Phe156 of loop β7-α6. The bound GMP was stabilized mainly through interactions of the guanine ring, which was sandwiched between Ile131 and the glycine-rich loop β3-α2 forming a network of hydrogen bonds to Lys44, Val49, Ala50 and Ala38. No contribution to binding was observed from the ribose and phosphate groups that were positioned outside the binding pocket in the solvent region.

For β-NAD and β-NADP, the observed electron density suggested that the entities of the bound ligands were likely nicotinamide-cleaved products, resulting in the complex of ADP-ribose and ADP-ribose-2’-phosphate (ADPRP), respectively. Although hydrolase activity catalyzing NAD hydrolysis^25^ has been documented for some macrodomains, we assumed that these cleavage products were likely a consequence of autohydrolysis of the compounds since no significant increase in ADP-ribose or ADPRP traces was observed in a HPLC assay upon an incubation of β-NAD or β-NADP with SARS-CoV-2 macrodomain (data not shown). Structural comparison demonstrated that both the NAD-cleaved product ADP-ribose and the NADP-cleaved product ADPRP assumed the binding mode highly identical to ADP-ribose (Figure 2 and 3D). Nevertheless, a difference was noted for ADPRP that the 2’-phosphate group, which was located in the solvent region, could engage additional magnesium-mediated contacts with an inward side chain of Asp157 (Figure 3D).

The flexibility of the ADP-ribose binding pocket with its ability to accept various ligands prompted us to speculate potential binding of anti-viral nucleoside analogues. We therefore selected a set of diverse clinically used antivirals, including abacavir, entecavir, acyclovir, gemcitabine, remdesivir and remdesivir metabolite GS-441524, and tested binding of these drugs using co-crystallization. Among the set, examination of electron density maps revealed that the remdesivir metabolite GS-441524 was the only ligand that showed binding in the crystal structures (2.15-Å resolution; Figure 4A and 4B). The interaction of this compound was rather unexpected as this metabolite and its active tri-phosphorylated form have been designed to target RNA-dependent RNA polymerase (RdRp) of several viruses, including Ebola virus, MERS-CoV, SARS-CoV and potentially SARS-CoV-2^26-29^. In addition, recent study has further demonstrated that the direct use of GS-441524 can potently inhibit the replication of SAR-CoV-2 in mouse model^30^. Based on our crystal structure, structural comparison demonstrated that the binding mode of GS-441524 in the macrodomain highly resembled that of the adenosine moiety of ADP-ribose. The amine group of the pyrrolotriazine ring, which was sandwiched between Val49 and Phe159, formed a hydrogen bond to Asp22 while the hydroxyl groups of the ribose moiety engaged two contacts with the backbone carbonyl atoms of Leu126 and Val155 (Figure 4C). The 1’-nitrile group was accommodated in the space adjacent to loop β7-α6. However, with a distance of ∼3.3 Å direct contacts to the backbone amine atoms of Phe156 and Asp157 might be weak, yet charge compatibility with this binding pocket was provided by the negative electrostatic potential of the nitrile group.

**Figure 4.**
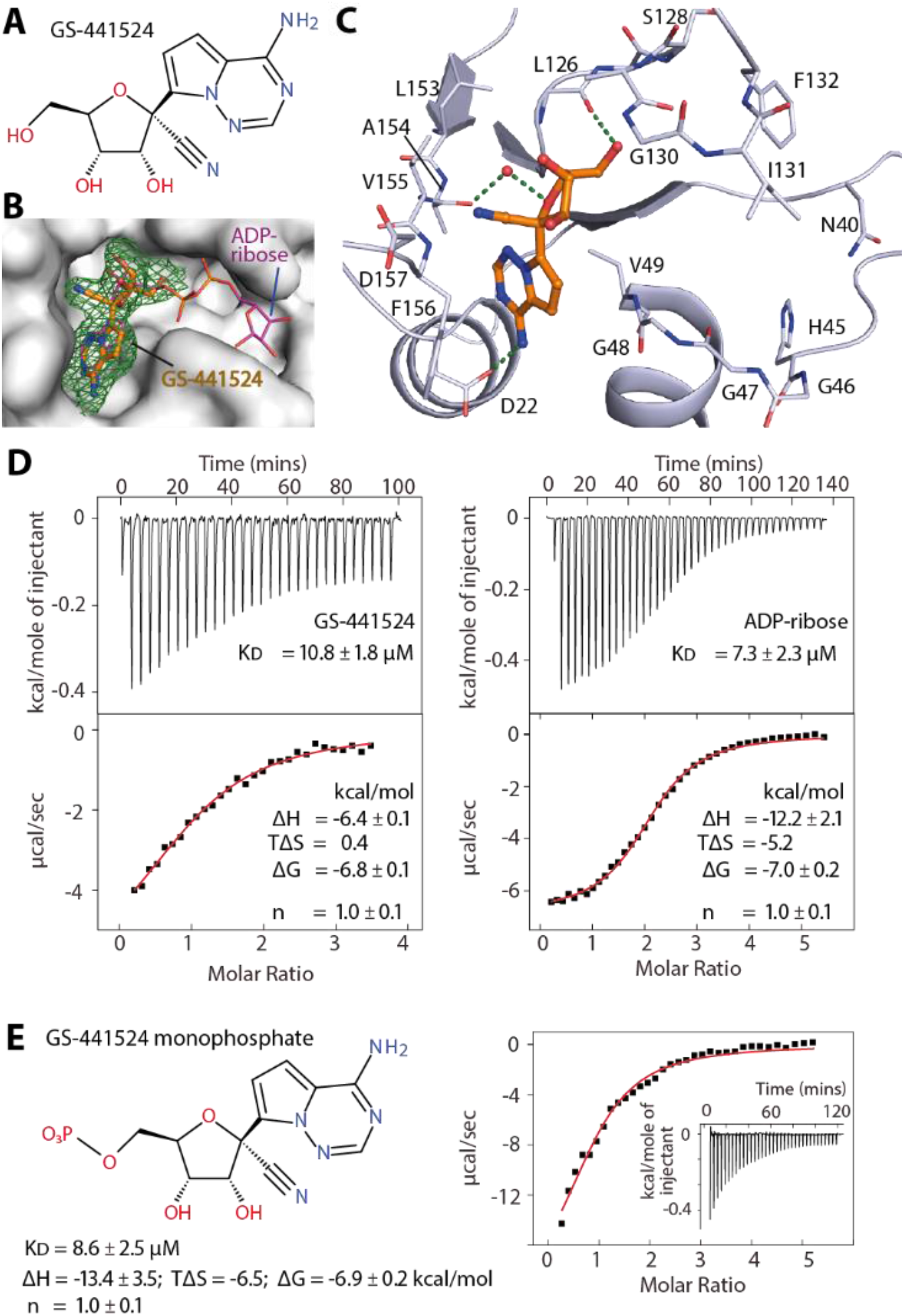
Interaction of the remdesivir metabolite GS-441524 with SARS-CoV-2 macrodomain. A) chemical structure of GS-441524. B) Surface representation revealing that GS-441524 occupied the binding site of the ADP-ribose adenosine moiety (pdb id 7bf6). The green mesh represents │*F*O│-│*F*C│ omitted electron density map contoured at 3σ for the bound ligand. C) Detailed interactions between the ligand and the protein. D) The ITC binding isotherm (top) and integrated heat of binding (bottom) for the interactions between either GS-441524 or ADP-ribose and the protein averaged from duplicates. E) Chemical structure of GS-441524 monophosphate and its ITC binding parameters from duplicate experiment for the maco-domain.

In comparison with the adenosine substrate, GS-441524 showed improved shape complementarity with the binding pocket (Figure 3A and 4C). To assess this, we performed isothermal calorimetry (ITC) to determine the affinities of this ligand, adenosine, AMP and ADP-ribose. Unfortunately, adenosine and AMP did not yield interpretable ITC binding isotherms. Nonetheless, we observed that the KDs of GS-441524 and ADP-ribose were remarkably comparable (10.8 and 7.3 μM, respectively). This was a surprising result considering that the two phosphate moieties and the terminal ribose, are missing in GS-441524. Thus, we synthesized the 5’ monophosphate derivative of this compound (GS-441524 monophosphate), and characterized the binding. Interestingly, an installation of 5’-phosphate led to a slight improved affinity with similar binding strength than ADP-ribose (KD of 8.6 μM), suggesting a contribution of the phosphate group for binding. Nevertheless, of particular note was different thermodynamics of the three ligands. The presence of phosphate group in GS-441524 monophosphate and ADP-ribose as well as additional terminal ribose-1’-phosphate moiety in the latter resulted in large negative binding enthalpy changes, which were counteracted by highly unfavorable entropy changes (TΔS) likely attributable to the constrained ligand and potentially pocket in the bound state.

To conclude, targeting the macrodomain offers an attractive target for the development of antiviral agents against SARS-CoV-2 and other viruses. To assist the inhibitor discovery efforts, we report a collective set of crystal structures extending our structural knowledge of this protein. Structural comparison demonstrated that the ADP-ribose binding site, essentially the pocket for ribose-1-phosphate, of the macrodomain possessed high structural plasticity. This was demonstrated by its adaptability to diverse naturally-occurring nucleosides and nucleotides including ADP-ribose-phosphate (ADPRP). This flexibility offers opportunities for rational targeting this protein module by small molecule inhibitors. Indeed, recent study has reported a number of potential small molecule binders discovered through crystallographic fragment screening^31^. In line with this, our unprecedented discovery of the binding of GS-441524, an active metabolite of remdesivir, supported this hypothesis that the ADP-ribose pocket of the macrodomain indeed represents a druggable site. The low micromolar affinity of GS-441524 and its small molecular weight offers a versatile starting point for ligand design.

## METHODS

### Protein production

The DNA encoding SARS-CoV-2 macrodomain was commercially synthesized, and subcloned into pET-28a(+) vector (Genscript, supplementary table s1). Expression of the recombinant protein harbouring an N-terminal His6 tags was carried out in *E. coli* Rosetta which was cultured in TB media. The culture was grown at 37 °C until reaching an OD600 of 1.6 prior to cooling to 18 °C. At an OD600 of 2.6-2.8, the protein expression was induced by adding 0.5 μM IPTG and the expression was continued overnight. Cells were harvested by centrifugation, and resuspended in buffer containing 50 mM Tris, pH 8.0, 500 mM NaCl, 10 mM imidazole, pH 7, 5% glycerol and 1 mM TCEP. Lysis was performed by sonication, and supernatant was clarified by centrifugation. The recombinant protein was initially purified using Ni^2+^-affinity chromatography, and subsequent cleavage of the histidine tag was performed by TEV protease treatment. The cleaved protein was passed through Ni^2+^-NTA column, and was further purified by size exclusion chromatography using Superdex S75. The pure protein was stored at -80 °C in buffer 25 mM Tris pH 8.0, 150mM NaCl and 5% glycerol.

### Crystallization and structure determination

The macrodomain was buffer exchanged into 25 mM Tris pH 8.0, 150mM NaCl and concentrated to ∼8.5-10 mg/ml. Crystallization was performed using sitting drop vapor diffusion method at 20 °C and conditions listed in supplementary table s2. Macroseeding was employed in all crystallization experiment using initial crystals of the macrodomain, which were stored in 30% broad-molecular-weight PEG smears^32^, M MgCl2, 0.1 M tris pH 7.0. For co-crystallization, the protein was mixed with 10-fold molar excess of the ligands. In the case of ADP-ribose, soaking was performed overnight using the ligand at 10 mM concentration. Viable crystals were cryo-protected with mother liquor supplemented with 20% ethylene glycol prior to flash cooling in liquid nitrogen.

Diffraction data were collected at Swiss Light Source, and were processed and scaled with XDS^33^ and aimless^34^, respectively. Initial structure solution was obtained with molecular replacement method using Phaser^35^ and the coordinate of the macrodomain (pdb id: 6wen). Manual model rebuilding alternated with refinement was performed in COOT^36^ and REFMAC5^37^. The geometry of the final models was verified using MOLPROBITY^38^. Omitted electron density maps for the bound ligands are shown in supplementary figure s1. The summary of data collection and refinement statistics are in supplementary table s2.

### Synthesis of GS-441524 monophosphate

Chemical synthetic procedure is described in supplementary information.

### Isothermal calorimetry

Isothermal titration calorimetry (ITC) experiment was performed using NanoITC instrument (TA Instrument) at 25 °C in the buffer containing 20 mM Tris pH 8.0, 100 mM NaCl, 0.5 mM TCEP, 5% glycerol. For ADP-ribose, the ligand at 500 µM in syringe was titrated into the reaction cell containing the protein at 106 µM, whereas for GS-441524 and GS-441524 monophosphate the protein at 250 µM was titrated into the reaction cell containing the compound at 22 μM. Data analyses were performed with NanoAnalyze software (TA Instrument) using an independent binding model from which the affinities and thermodynamic parameters were calculated.

## AUTHOR INFORMATION

## Author Contributions

‡ X.N. and M.S. contributed equally. All authors have given approval to the final version of the manuscript.

## ABBREVIATIONS

nsP3: non-structural protein 3
MD: macrodomain
ADP-ribose: adenosine diphosphate ribose
AMP: adenosine monophosphate
GMP: guanosine monophosphate
NAD: nicotinamide adenine dinucleotide
NADP: nicotinamide adenine dinucleotide phosphate
ADPRP: adenosine diphosphate ribose phosphate

