## supplementary table for "Structural insights into plasticity and discovery of remdesivir metabolite GS-441524 binding in SARS-CoV-2 macrodomain"

### Supplementary information


| | Page |
|---|---|
| **Supplementary figure s1.** $\|F_O\|-\|F_C\|$ omitted electron density map contoured at 3σ for the bound ligands. | S2 |
| **Supplementary table s1.** Details of recombinant SARS-CoV-2 macrodomain. | S3 |
| **Supplementary Table s2.** Data collection and refinement statistics. | S4 |
| **Supplementary method.** Synthesis of GS-441524 monophosphate | S6 |

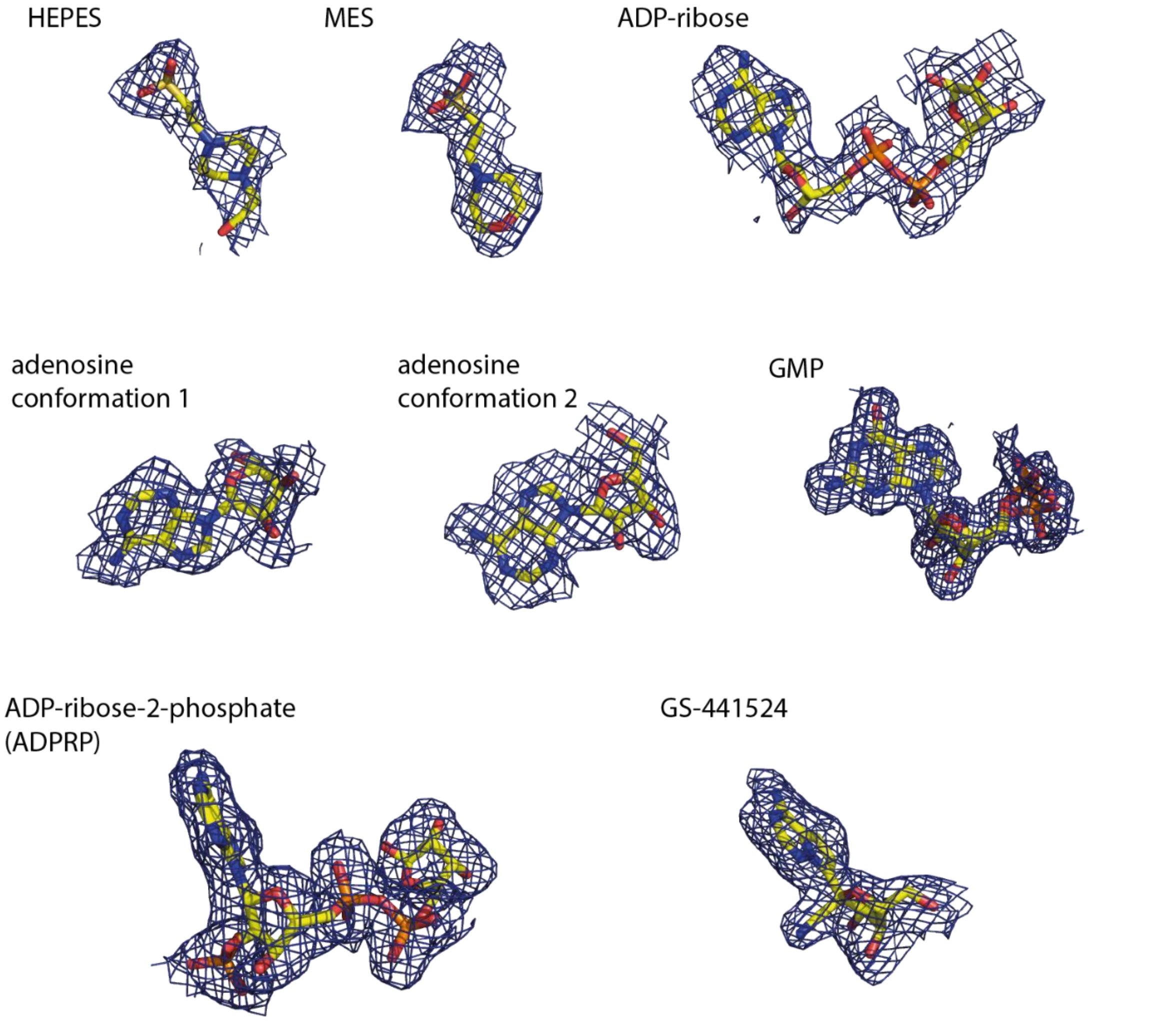


**Supplementary Figure s1.** $|F_O|-|F_C|$ omitted electron density map contoured at 3σ for the bound ligands.

**Supplementary table s1.** Details of recombinant SARS-CoV-2 macrodomain.

| | Vector | Recombinant protein sequence |
|---|---|---|
| **SAR-CoV-2 macrodomain** | **pET-28a(+)** | **MGSSHHHHHHSSGENLYFQGHMVNSFSGYLKLTDNVYIKNADIVEEAKKVKPTVVVNAANVYLKHGGGVAGALNKATNNAMQVESDDYIATNGPLKVGGSCVLSGHNLAKHCLHVVGPNVNKGEDIQLLKSAYENFNQHEVLLAPLLSAGIFGADPIHSLRVCVDTVRTNVYLAVFDKNLYDKLVSSFLEMK** |

**Supplementary Table s2.** Data collection and refinement statistics.

| **Complex** | **apo/HEPES** | **apo/MES** | **ADP-ribose** |
|---|---|---|---|
| PDB codes | **6ywk** | **6ywm** | **6ywl** |
| Beamline | SLS X06SA | SLS X06SA | SLS X06SA |
| *Data Collection* | | | |
| Resolution[a] (Å) | 49.09-2.20 (2.28-2.20) | 49.22-2.16 (2.24-2.16) | 48.83-2.50 (2.64-2.50) |
| Space group | $P\ 2_12_12_1$ | $P\ 2_12_12_1$ | $P\ 2_12_12_1$ |
| Cell dimensions | a=39.2, b=111.8, c=196.4 Å<br>α=β=γ=90.0 | a=37.8, b=109.1, c=114.4 Å<br>α=β=γ=90.0 | a=38.4, b=111.9, c=195.3 Å<br>α=β=γ=90.0° |
| Number of unique reflections[a] | 45,087 (4,348) | 26,281 (2,558) | 30,002 (4,288) |
| Completeness[a] (%) | 100.0 (99.9) | 100.0 (100.0) | 99.4 (99.3) |
| I/σI[a] | 10.7 (2.0) | 8.3 (2.0) | 6.9 (1.9) |
| $R_{merge}$[a] (%) | 0.138 (0.925) | 0.162 (0.873) | 0.199 (0.930) |
| CC (1/2) [a] | 0.998 (0.762) | 0.995 (0.787) | 0.990 (0.736) |
| Redundancy[a] | 8.5 (7.9) | 6.7 (6.9) | 5.6 (5.9) |
| *Refinement* | | | |
| Number atoms in refinement (P/L/O)[b] | 6,496/ 15/ 424 | 3,885/ 24/ 287 | 6,472/ 180/ 248 |
| B factor (P/L/O)[b] (Å$^2$) | 39/ 76/ 48 | 34/ 57/ 39 | 40/ 31/ 38 |
| $R_{fact}$ (%) | 17.6 | 17.5 | 18.9 |
| $R_{free}$ (%) | 21.4 | 22.9 | 22.3 |
| rmsd bond[c] (Å) | 0.013 | 0.013 | 0.010 |
| rmsd angle[c] (°) | 1.4 | 1.3 | 1.1 |
| *Molprobity Ramachandran* | | | |
| Favor (%) | 99.65 | 99.01 | 98.11 |
| Outlier (%) | 0 | 0 | 0 |
| Crystallization condition | 33% broad-molecular-weight PEG smears, 0.1 M $MgCl_2$, 0.1 M HEPES, pH 7.0 | 23% PEG 6000, 0.1 M $MgCl_2$, 5% ethylene glycol, 0.1 M MES, pH 6.0 | 27% PEG 4000, 0.2 M sodium acetate, 0.05 M $MgCl_2$, 0.1 M tris, pH 8.0 |

[a] Value in brackets indicates high-resolution shell statistics.
[b] P/L/O indicates protein, ligands and others.
[c] rmsd indicates root-mean-squre deviation.

**Supplementary Table s2.** ***(continued)*** Data collection and refinement statistics.

| Complex | Adenosine | GMP | ADPRP | GS-441524 |
|---|---|---|---|---|
| PDB codes | **7bf3** | **7bf4** | **7bf5** | **7bf6** |
| Beamline | SLS X06SA | SLS X06DA | SLS X06SA | SLS X06SA |
| *Data Collection* | | | | |
| Resolution[a] (Å) | 48.99-2.00 (2.07-2.00) | 36.26-1.55 (1.60-1.55) | 48.80-2.05 (2.12–2.05) | 48.43-2.15 (2.23–2.15) |
| Space group | $P\ 2_12_12_1$ | $P\ 4_1$ | $P\ 2_12_12_1$ | $C\ 2$ |
| Cell dimensions | a=39.2, b=111.4, c=196.0 Å | a=b=72.5, c=33.4 Å | a=38.6, b=111.3, c=195,2 Å | a=157.2, b=30.5, c=111,7 Å |
| | $\alpha$=β=γ=90.0 | $\alpha$=β=γ=90.0 | $\alpha$=β=γ=90.0° | $\alpha$= γ=90.0°, β=119.9° |
| Number of unique reflections[a] | 59,412 (5,774) | 25,315 (2,287) | 53,878 (5,196) | 25,440 (2,459) |
| Completeness[a] (%) | 100.0 (100.0) | 99.2 (92.8) | 99.7 (99.8) | 99.1 (99.2) |
| I/σI[a] | 10.9 (2.0) | 13.8 (2.6) | 8.3 (1.9) | 10.7 (1.9) |
| $R_{merge}$[a] (%) | 0.127 (0.885) | 0.064 (0.349) | 0.141 (0.839) | 0.090 (0.755) |
| CC (1/2)[a] | 0.998 (0.735) | 0.998 (0.807) | 0.995 (0.695) | 0.998 (0.677) |
| Redundancy[a] | 7.5 (7.5) | 6.1 (3.3) | 6.2 (6.3) | 5.3 (5.2) |
| *Refinement* | | | | |
| Number atoms in refinement (P/L/O)[b] | 6,538/ 38/ 628 | 1,329/ 48/ 241 | 6,500/ 160/ 569 | 3,838/ 63/ 168 |
| B factor (P/L/O)[b] (Å$^2$) | 29/ 60/ 37 | 14/ 13/ 30 | 28/ 41/ 35 | 49/ 39/ 43 |
| $R_{fact}$ (%) | 17.5 | 13.8 | 17.7 | 18.0 |
| $R_{free}$ (%) | 21.7 | 17.4 | 21.7 | 22.6 |
| rmsd bond[c] (Å) | 0.013 | 0.018 | 0.014 | 0.012 |
| rmsd angle[c] (°) | 1.4 | 1.7 | 1.4 | 1.4 |
| *Molprobity* | | | | |
| *Ramachandran* | | | | |
| Favor (%) | 98.94 | 99.40 | 97.64 | 99.00 |
| Outlier (%) | 0 | 0 | 0 | 0 |
| Crystallization condition | 33% broad-molecular-weight PEG smears, 0.1 M $MgCl_2$, 0.1 M tris, pH 7.0 | 30% PEG 4000, 0.2 M sodium acetate, 0.1 $MgCl_2$, 0.1 M tris, pH 8.3 | 30% broad-molecular-weight PEG smears, 0.1 M $MgCl_2$, 0.1 M tris, pH 7.0 | 30% PEG 4000, 0.2 M sodium acetate, 0.1 M tris, pH 8.3 |

[a] Value in brackets indicates high-resolution shell statistics.

[b] P/L/O indicates protein, ligands and others.

[c] rmsd indicates root-mean-squre deviation.

**Supplementary method.** Synthesis of GS-441524 monophosphate

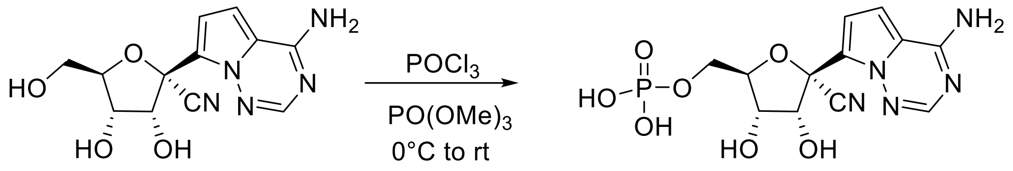


**GS-441524 monophosphate**

A solution of GS-441524 (43.7 mg, 0.15 mmol) in trimethyl phosphate (1.5 mL) was stirred in a sealed tube under Ar at rt for 15 min. The solution was then cooled to 0°C and freshly distilled phosphorous oxychloride (21.2 µL, 0.225 mmol) was added dropwise. The resulting solution was stirred at rt for 1h. Further 100 µL of phosphorous oxychloride were added at rt and the resulting solution was stirred at rt for 1h (full conversion by HPLC). The reaction mixture was quenched with water at 0°C and directly purified by preparative HPLC to obtain 43.6 mg (78%) of the expected product as a white solid. $^{1}H$ NMR (300 MHz, $D_2O$) δ 7.98 (s, 1H), 7.27 (d, *J* = 4.9 Hz, 1H), 7.05 (d, *J* = 4.9 Hz, 1H), 4.83 (d, *J* = 5.2 Hz, 1H), 4.43-4.39 (m, 1H), 4.31 (t, *J* = 4.7 Hz, 1H), 4.04-3.92 (m, 2H); $R_f$ HPLC: 3.4 Min (13 Min from 10 to 95% MeCN in water (0.1 % formic acid), then 7 min 95% MeCN). 95.7 % purity; HRMS (MALDI): m/z found. 372.0705 $[M+H]^+$ (cal. $C_{12}H_{15}N_5O_7P$ 372.0704).

To record NMR-spectra, the compound was dissolved in $D_2O$ and measured on Avance 300 from Bruker Corporation (Massachusetts, USA). All chemical shift values are reported in ppm, the multiplicity of the signals assigned as follows: s (singlet), d (duplet), t (triplet) and m (multiplet). Mass spectrometry analysis was performed in positive ion mode by electrospray-ionization (ESI) on a LCMS-2020 single quadrupole MS from Shimadzu (Duisburg, Deutschland). Precision mass was measured using MALDI Orbitrap XL from Life Technologies GmbH (Darmstadt, Germany). For purity estimation of the synthesized compounds, a reverse phase high-performance liquid chromatography (RP-HPLC) was performed using the Luna 10 µm C18(2) 100 Å, LC Column 250 x 4.6 mm from Phenomenex LTD (Aschaffenburg, Germany) and the analysis was conducted using the Shimadzu prominence module from Shimadzu. Acetonitrile and aqueous formic acid 0.1% were used as eluents. The established method for purity determination was initiated with 90% water (0.1% formic acid), then a linear gradient from 90% to 5% water (0.1% formic acid) for 13 min was chosen, finally additional 7 min 5% water (0.1% formic acid). The flow rate was adjusted to 1.0 mL/min and the UV–vis detection occurred at 254 nm and 280 nm, respectively.
